# A synNotch-based morphogen detection system reveals sFRP2 enhances Wnt3a signaling

**DOI:** 10.64898/2026.02.09.704138

**Authors:** Kosuke Mizuno, Satoshi Toda

**Author notes:** Correspondence address: Satoshi Toda, PhD, Institute for Protein Research, The University of Osaka, Suita, Japan.

## Abstract

Morphogen gradients provide essential positional information during tissue development, yet the extracellular mechanisms that regulate morphogen transport and presentation remain poorly understood. Here, we introduce a mechanosensitive detection system based on synthetic Notch (synNotch) receptors that selectively detects surface-bound, but not soluble, morphogen complexes. Applying this platform to Wnt signaling, we demonstrate that secreted Frizzled-related protein 2 (sFRP2) promotes the recruitment of Wnt3a to the cell surface via heparan sulfate proteoglycans, enabling coordinated endocytosis and robust activation of canonical Wnt/β-catenin signaling. Notably, sFRP2 extends the effective signaling range of Wnt3a and amplifies Wnt responses under ligand-limiting conditions. In intestinal organoid cultures, sFRP2 enhances Wnt3a-driven growth and induces high-Wnt morphological states with low-level Wnt concentrations. These findings identify sFRP2 as an extracellular carrier that stabilizes surface-bound Wnt3a and regulates both the strength and spatial range of Wnt signaling. More broadly, this work demonstrates the utility of synNotch mechanosensing for dissecting extracellular morphogen dynamics and highlights morphogen carrier proteins as a platform for optimizing organoid culture.

## Introduction

Morphogens, such as Wnt, Sonic hedgehog (Shh), bone morphogenetic proteins (BMPs), and retinoic acid (RA), are secreted signaling molecules that play fundamental roles in orchestrating complex biological processes.^1^ They diffuse through tissues to establish concentration gradients that provide essential positional information to receiving cells, which determine their specific cell fates. Morphogen-mediated regulation can be broadly divided into two distinct phases. The first phase is the extracellular diffusion and delivery of the morphogen to target cells. The second phase involves the subsequent cellular responses and the resulting changes in the properties of the receiving cells. To date, the second phase has been extensively characterized in various developmental contexts. For instance, studies using both in vitro and in vivo models have demonstrated that transcriptional networks and cell adhesion play crucial roles in forming stripe tissue patterns in response to morphogen gradients.^2–7^ In contrast, due to technical difficulties in visualizing diffusing morphogens inside developing tissues, the physical and biochemical processes that govern the transport of morphogens from their source to distant target cells remain largely elusive.

Among the various families of morphogens, Wnt proteins represent prototypical examples of molecules that require intricate regulation in the extracellular space due to their high hydrophobicity.^8,9^ Wnt proteins undergo post-translational modification with palmitoleic acid at a conserved serine residue.^9^ To solubilize and stabilize Wnt in the extracellular environment, this lipid moiety must be shielded through the formation of specific complexes. These include homo-oligomers or heterodimers with carrier proteins, such as the serum glycoprotein afamin and secreted Frizzled-related proteins (sFRPs).^10–12^ During their transport to the cell surface, Wnt molecules and Wnt3a/sFRP2 complexes are known to associate with heparan sulfate proteoglycans (HSPGs).^13–16^ To investigate the extracellular dynamics of Wnt proteins, recent studies using a functional fluorescent-tagged Wnt3a in analytical ultracentrifugation coupled with a fluorescence detection system (AUC-FDS) have elucidated the composition of Wnt3a complexes in culture media, revealing that Wnt3a does not exist as a monomer but instead forms homo-trimers as its minimum unit or preferentially forms heterodimers with afamin (**Fig. 1A**).^17^ However, the absence of simple assays to quantify Wnt3a-cell surface interactions within complex extracellular environments has limited our understanding of which specific complexes preferentially associate with the cell surface and how they contribute to the activation of intracellular signaling.

**Figure 1.**
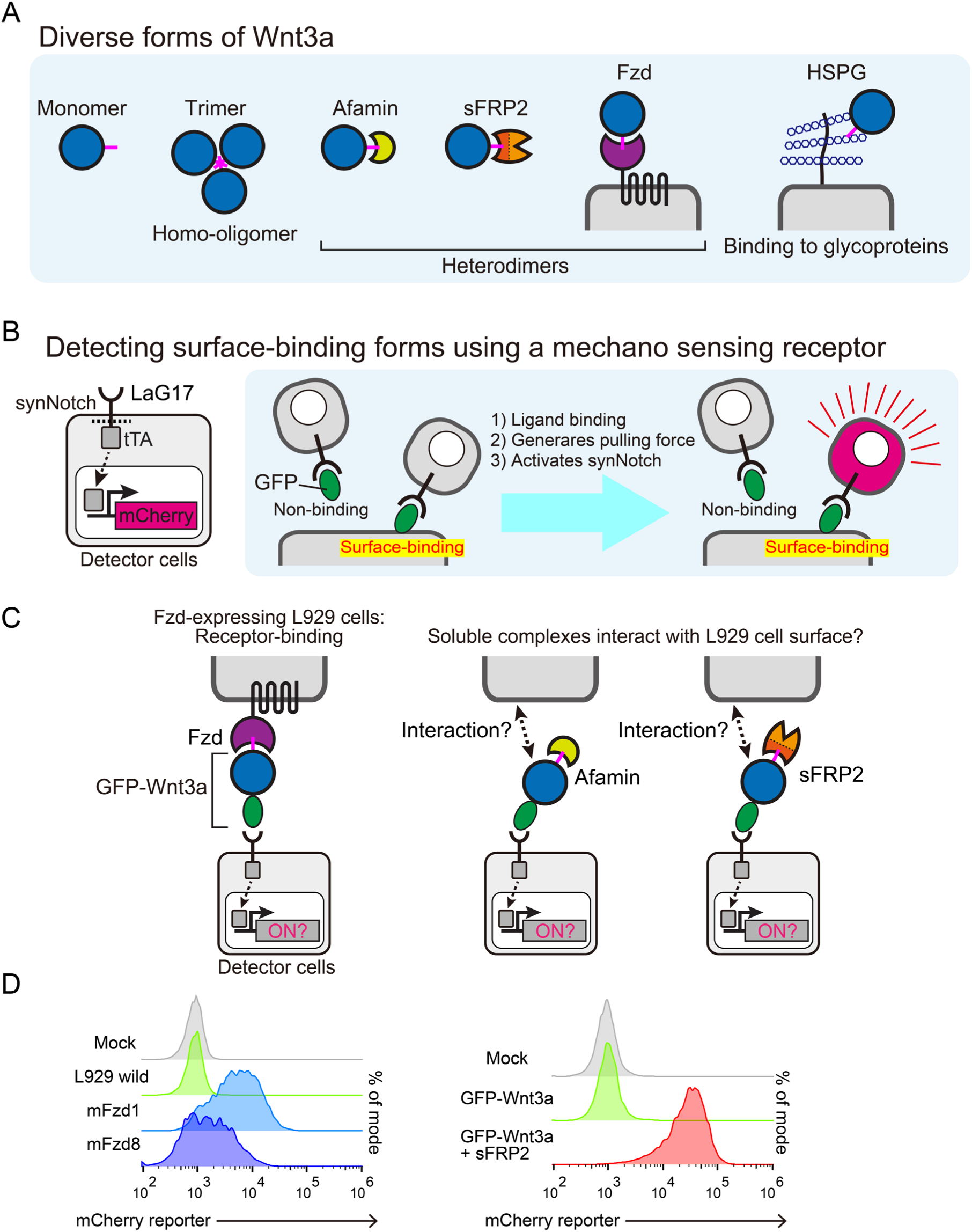
Exploring Wnt3a-regulatory factors based on synthetic Notch mechano-sensing receptor. (A) Extracellular diversity of Wnt3a conformations. Schematic representation of the various reported forms of Wnt3a present in the extracellular environment. (B) Detection mechanism for surface-bound proteins using synNotch receptor. Detector cells are engineered to express the anti-GFP nanobody (LaG17)-tTA synNotch receptor and a responsive mCherry reporter cassette. Mechanical pulling force is required to trigger receptor proteolysis and subsequent tTA release. Consequently, the system specifically responds to surface-bound GFP ligands but not unbound soluble GFP in the medium. (C) Experimental setup for monitoring Wnt3a recruitment to the cell surface. Detector cells were cocultured with L929 cells expressing Wnt3a receptors, such as Fzd1 or Fzd8 in conditioned media containing GFP-Wnt3a. GFP-Wnt3a forms a complex with afamin derived from serum in the medium.^17^ In another experiment, detector cells alone were cultured with conditioned media containing GFP-Wnt3a and sFRP2. (D) After 24-hour incubation, the mCherry fluorescence of detector cell were measured by flow cytometry. Coculturing with Fzd-expressing cells resulted in reporter activation, indicating that GFP-Wnt3a alone or GFP-Wnt3a/afamin complexes do not interact with the cell surface at a level sufficient to trigger the synNotch receptor (left). The addition of sFRP2 induced a robust increase in mCherry signal, demonstrating that sFRP2 effectively recruits Wnt3a to the cell surface (right).

Over the past decade, various synthetic biology approaches using artificial receptors have been developed to manipulate intercellular communication.^18–22^ These tools have been instrumental in exploring the minimal requirements for morphogenesis and in advancing cancer immunotherapies, such as chimeric antigen receptor (CAR)-T cell therapies.^23–25^ In particular, synthetic Notch (synNotch) receptor converts ligand-binding inputs into customized transcriptional outputs.^19^ The synNotch activation requires a mechanical pulling force between the ligand and the extracellular domain of the receptor.^26,27^ This unique mechanism implies that the ligand must be immobilized on a surface, including the plasma membrane of adjacent cells, the extracellular matrix (ECM), or a culture plate.^19,28,29^

Based on this mechanical requirement of synNotch system, we hypothesized that synNotch could serve as a powerful platform for selectively detecting molecules bound to the cell surface (**Fig. 1B**). The activation status of synNotch system directly indicates whether target molecules associate with the cell surface or not. Furthermore, this system is particularly advantageous for detecting weak interactions that are challenging to visualize by conventional microscopy, as ligand-binding events are amplified into reporter gene expression. This enables a more efficient and quantitative evaluation of surface binding by flow cytometry. In this study, we employed the synNotch system as a highly sensitive detector for surface-bound Wnt3a complexes. Using GFP-Wnt3a, we addressed the fundamental question of which specific Wnt3a complexes preferentially associate with the cell surface. Our findings demonstrate that Wnt3a/sFRP2 complexes strongly interact with the cell membrane and significantly enhance Wnt/β-catenin signaling. Furthermore, our analysis using intestinal organoids reveals that sFRP2 promotes Wnt3a activity in a physiologically relevant context.

## Results

### synNotch-based system for detecting surface-bound Wnt3a complexes

To specifically detect surface-bound Wnt3a complexes among various forms of Wnt3a complexes, we established a synNotch-based detector cell line. This detector cell line, derived from the mouse fibroblast line L929, expresses synNotch receptors bearing an extracellular anti-GFP nanobody (LaG17). Upon recognition of GFP-tagged ligands, the transmembrane domain of synNotch receptor undergoes proteolytic cleavage, resulting in nuclear translocation of the intracellular domain and subsequent induction of the red fluorescent protein mCherry as a reporter (**Fig. 1B**).

We first validated the system by testing whether the detector cells could sense GFP-Wnt3a immobilized on the surface of adjacent cells. The detector cells were co-cultured with L929 cells overexpressing Wnt3a receptors (Frizzleds, Fzd) in the presence of GFP-Wnt3a-containing conditioned medium (GFP-Wnt3a CM) (**Fig. 1C**).^17^ As shown in **Figure 1D**, robust activation of detector cells was observed when GFP-Wnt3a was captured on cells overexpressing mouse Fzd1 or Fzd8.

We next investigated the behavior of Wnt3a complexes in the presence of carrier proteins. In standard culture media, Wnt3a predominantly exists as a soluble heterodimer with serum-derived protein afamin.^10^ Consistent with this, no detectable activation of the detector cells was observed with GFP-Wnt3a CM alone, suggesting that soluble Afamin/Wnt3a complexes do not associate with the cell surface. We then examined Wnt3a complexes formed in the presence of sFRP2. Addition of sFRP2 alters the composition of Wnt3a complexes toward the formation of Wnt3a/sFRP2 complexes.^17^ Notably, co-incubation with GFP-Wnt3a CM and sFRP2-containing conditioned medium (sFRP2 CM) led to strong activation of the detector cells (**Fig. 1D**). This result is consistent with recent reports showing that the Wnt3a/sFRP2 complexes preferentially interact with the cell surface, thereby facilitating the release of Wnt3a-containing exosomes.^16^ Together, these results indicate that the synNotch-based detector systems can successfully distinguish soluble and surface-bound forms of Wnt3a complexes, although the precise molecular mechanisms underlying Wnt3a/sFRP2-cell surface interactions and their physiological significance remain to be elucidated.

### sFRP2 acts as a molecular scaffold for anchoring Wnt3a to the plasma membrane across multiple cell lines

To determine whether synNotch-based detection of Wnt3a complexes is specific to L929 fibroblasts or can be broadly applied to different types of cells, we established detector cell lines using epithelial MDCKII and EpH4, as well as another fibroblast line NIH3T3. In all tested cell lines, the detector system consistently detected surface-bound Wnt3a complexes, demonstrating the robustness and general applicability of this approach (**Fig. 2A**). Fluorescence microscopy imaging of L929 detector cells confirmed induction of the mCherry reporter by GFP-Wnt3a in the presence of sFRP2, accompanied by increased uptake of GFP-Wnt3a following synNotch activation (**Fig. 2B**).

**Figure 2.**
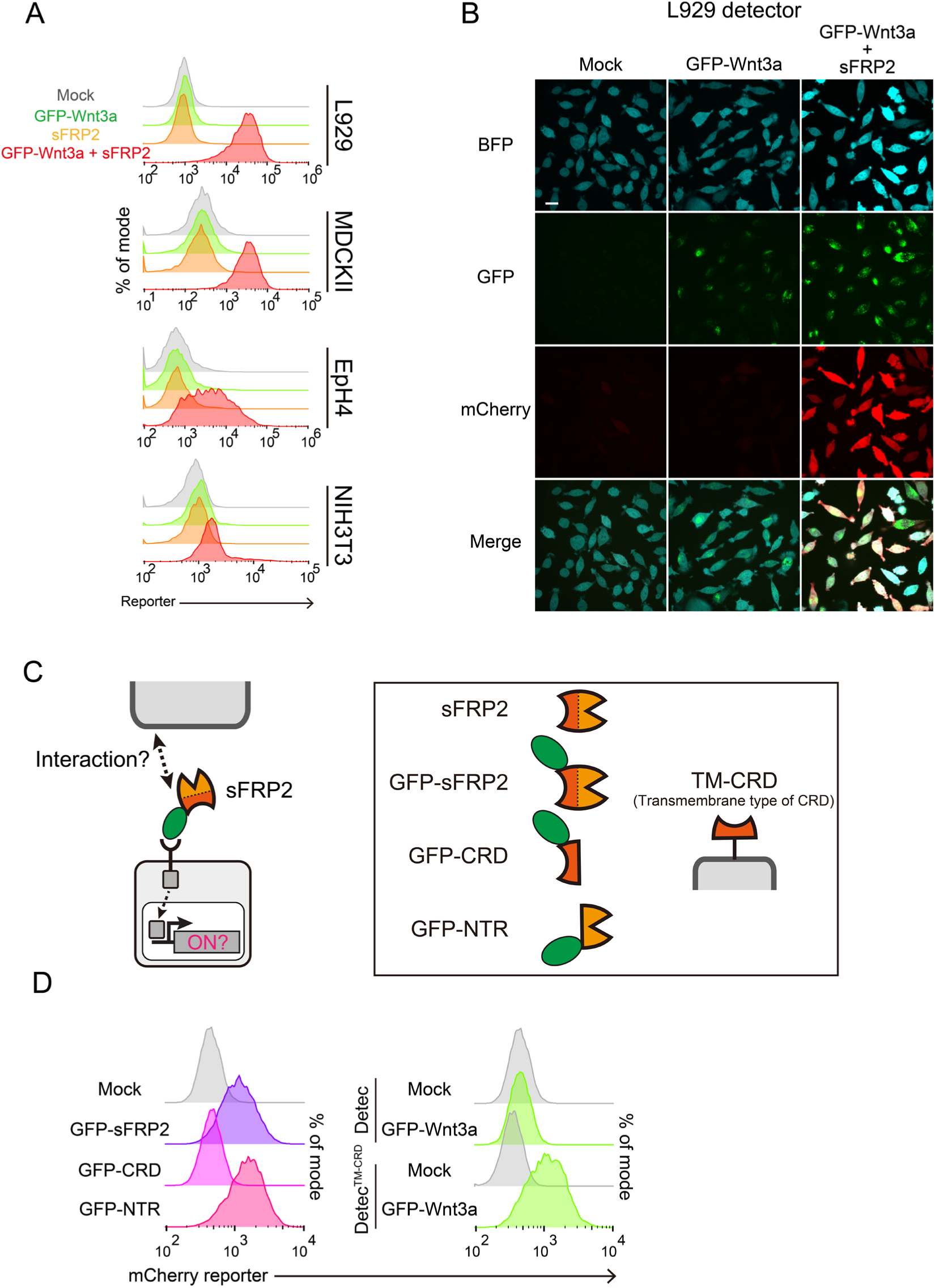
Characterization of sFRP2 as a cell surface binder of Wnt3a. (A) Activation of detector cells in multiple cell lines. Receptor functionality was confirmed in L929, MDCKII, EpH4, and NIH3T3 cell lines. These cells express the anti-GFP (LaG17)-tTA synNotch receptor, with the exception of MDCKII cells, which express the anti-GFP (LaG2) variant. The reporter protein used was mCherry for L929, MDCKII, and EpH4, while tagBFP was utilized for NIH3T3 cells. After 24-hour incubation with conditioned media containing GFP-Wnt3a and sFRP2, reporter expression levels were quantified by flow cytometry. The detection system was functional in all tested cell lines upon treatment with the GFP-Wnt3a and sFRP2 supernatant. (B) Confocal images of L929 detector cells. The images were acquired immediately before FACS analysis after 24-h incubation. tagBFP is stably expressed in detector cells as a marker for reporter cassette integration. GFP internalization occurs accompanied with anti-GFP synNotch activation. mCherry expression indicates successful reporter activation. Scale bar: 20 µm. (C) Schematic representation of the chimeric sFRP2 proteins. GFP was fused to the N-terminus of specific sFRP2 domains. For transmembrane type of CRD (TM-CRD), the CRD of sFRP2 was fused to the transmembrane domain of PDGFRα. (D) Domain-specific requirements for surface recruitment. Flow cytometric analysis was performed using conditioned media containing indicated chimeric proteins. Detector cells were incubated with these supernatants for 24 hours. In the absence of GFP-Wnt3a, both GFP-sFRP2 and GFP-NTR induced reporter activation, suggesting that the NTR is essential for the cell-surface binding of sFRP2. Conversely, in the absence of sFRP2, GFP-Wnt3a alone was sufficient to activate detector cells expressing the membrane-tethered TM-CRD, indicating that CRD is essential for the interaction with Wnt3a.

To further investigate the molecular basis of Wnt3a/sFRP2 association with cell surface, we examined which domain of sFRP2 mediates this interaction. sFRP2 consists of an N-terminal cysteine-rich domain (CRD) and a C-terminal netrin-like domain (NTR). We therefore generated following constructs: GFP-sFRP2, GFP-CRD, GFP-NTR, and transmembrane-anchored CRD (TM-CRD) (**Fig. 2C and Fig. S1A**).

Using synNotch-based L929 detector cells, we assessed their ability to interact with the cell surface. Detector activation was observed with GFP-sFRP2 and GFP-NTR, but not with GFP-CRD (**Fig. 2D**). These results indicate that sFRP2 primarily associates with the cell surface via its NTR domain, consistent with previous reports regarding its affinity with the cell surface.^14,30^ Additionally, detector cells expressing TM-CRD were activated by GFP-Wnt3a, demonstrating that the CRD is sufficient to bind with Wnt3a and that membrane anchoring of the CRD enables efficient capture of GFP-Wnt3a from the extracellular environment onto cell surface.

To exclude the possibility that detector cells were activated by sFRP2 non-specifically adsorbed to the culture plate rather than bound to cell membranes, we performed experiments using a three-dimensional spheroid culture system (**Fig. S1B**). Under these 3D conditions in which detector cells did not attach to plate surface, robust mCherry expression was still observed, indicating that the detector activation was driven by interactions with sFRP2 molecules localized on the surfaces of neighboring cells (**Fig. S1C**).

We further examined the localization of GFP-Wnt3a/sFRP2 complexes and GFP-tagged sFRP2 mutants in MDCKII cells by confocal fluorescence microscopy (**Fig. S2A-B**). Incubation of MDCKII cells with GFP-Wnt3a CM resulted in both surface-associated and internalized GFP signals in the presence of sFRP2. Similarly, GFP-sFRP2 and GFP-NTR exhibited comparable localization patterns on cell surface and inside, confirming consistent behavior across cell types.

### HSPG-dependent membrane association and internalization of the Wnt3a/sFRP2 complex

Recent studies have reported that Wnt3a/sFRP2 complexes interact with heparan sulfate proteoglycans (HSPGs) on cell surface.^14,16^ To validate this interaction in our system, we generated detector cells expressing a membrane-tethered form of Heparinase III (HepIII), an enzyme that degrades heparan sulfate chains (**Fig. 3A**).^31^ Expression of HepIII in detector cells markedly reduced synNotch activation induced by GFP-Wnt3a and sFRP2 CMs (**Fig. 3B**). Similar inhibition was observed when recombinant sFRP2 was added at various concentrations, confirming that HSPG degradation substantially suppresses detector activation (**Fig. S3**). These results indicate that Wnt3a/sFRP2 complexes associate with the cell surface primarily through interaction with HSPGs in L929 cells.

**Figure 3.**
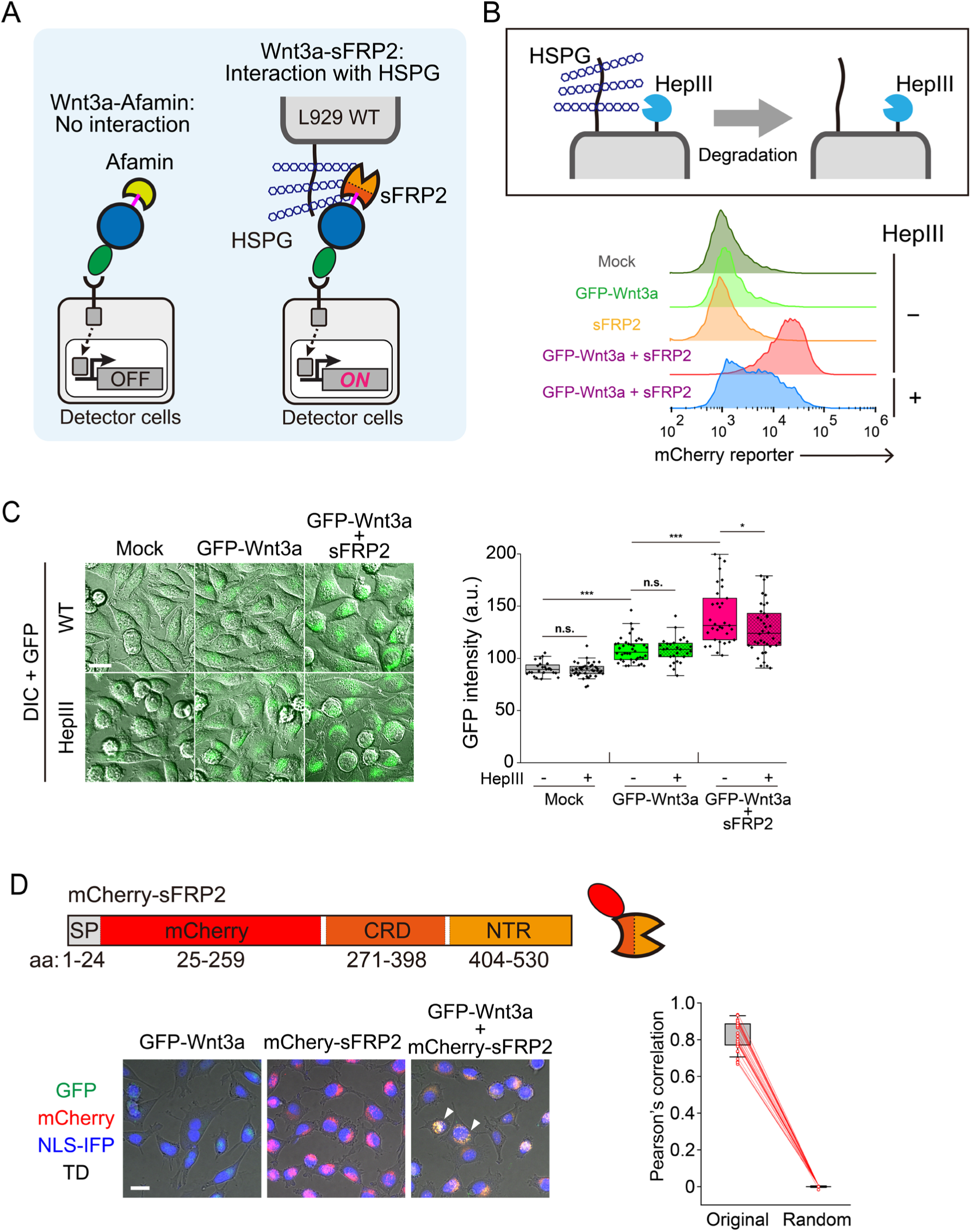
sFRP2 recruits Wnt3a to cell surface via HSPG and promotes their co-internalization into L929 cells. (A) Wnt3a/sFRP2 complexes interact with HSPG on the cell surface, while Afamin/Wnt3a shows no interaction. (B) Schematic illustration of the membrane-tethered HepIII system (top). HepIII-HA-GPI was stably expressed in detector cells and wild type L929 cells to enzymatically degrade cell-surface heparan sulfate. HepIII specifically cleaves the glycosidic bonds of heparin and heparan sulfate. Impact of heparan sulfate depletion on detector activation (bottom). mCherry induction levels were compared between standard detector cells and HepIII-expressing detector cells using flow cytometry. The mCherry expression was significantly reduced in HepIII-expressing cells, although it was not completely inhibited to basal levels. (C) Quantification of GFP-Wnt3a recruitment and internalization. After treating L929 cells with conditioned medium for 24 hours, confocal images were acquired for the GFP channel and differential interference contrast (DIC) (left). Scale bar: 20 µm. The total GFP intensity in each single cell was quantified after performing segmentation (right). Treatment with GFP-Wnt3a CM alone slightly increased GFP signals compared to the mock control, while the combined treatment with sFRP2 CM further enhanced these signals. HepIII expression resulted in a slight reduction of GFP recruitment specifically in the combined treatment group. Statistical significance was determined using Welch’s t-test with *** for P < 0.001, * for P < 0.05, and n.s. (non-significant) for P ≧ 0.05. (D) An mCherry-sFRP2 construct was generated by fusing mCherry to the N-terminus of sFRP2 to visualize the distribution of Wnt3a and sFRP2 at the same time. The intracellular localization of these proteins was monitored using confocal microscopy. Representative images display GFP-Wnt3a alone (left), mCherry-sFRP2 alone (center), and the combined treatment of GFP-Wnt3a and mCherry-sFRP2 (right). The colocalization of green and red signals results in yellow fluorescence, appearing as small puncta within the perinuclear region (white arrowheads). Scale bar: 20 μm. To quantitatively evaluate the interaction between GFP-Wnt3a and mCherry-sFRP2, each single cell was automatically segmented using CellPose2.0, then Pearson’s correlation coefficients were calculated. To exclude the possibility of coincidental overlap, the correlation values obtained from the original dual-channel images were compared against those derived from randomly shuffled GFP and mCherry channels.

We next analyzed the localization of GFP-Wnt3a and sFRP2 in L929 cells by confocal microscopy. Consistent with results using detector cell, total cellular GFP intensity, including both surface binding and internalized puncta, was significantly reduced in HepIII-expressing cells although not completely negated (**Fig. 3C**). This partial reduction suggests that HSPGs were not completely degraded by HepIII expression or that additional cell surface molecules may associate with Wnt3a/sFRP2 complex. A similar trend was observed for GFP-sFRP2 and GFP-NTR localization in L929 cells (**Fig. S4A**). In contrast, HepIII expression in MDCKII cells eliminated most of GFP signals derived from GFP-sFRP2 and GFP-NTR. These results indicate that cell type-specific membrane compositions influence the interaction between sFRP2 and cell surface (**Fig. S4B**).

Co-incubation of GFP-Wnt3a CM with L929 cells revealed that GFP-Wnt3a/sFRP2 complexes predominantly accumulated within the cytoplasmic vesicles rather than localizing on the cell membrane. In contrast, MDCKII cells exhibited clearer membrane localization, with GFP-positive intracellular vesicles also visible (**Fig. S2A-B**). These results indicate that endocytosis of Wnt3a/sFRP2 complexes occurs across multiple cell types.^16^ To determine whether Wnt3a and sFRP2 remain associated after internalization or sFRP2 proteins are trafficked independently, we generated mCherry-sFRP2 and simultaneously analyzed the localization of GFP-Wnt3a and mCherry-sFRP2. Colocalization analysis revealed that the majority of GFP and mCherry fluorescence signals overlapped significantly within intracellular puncta. Quantitative assessment yielded a high Pearson’s correlation coefficient, demonstrating that Wnt3a and sFRP2 are highly colocalized even after internalization (**Fig. 3D and Fig. S5**). These data indicate that Wnt3a/sFRP2 complexes remain intact following endocytosis and accumulate in intracellular vesicles.

### sFRP2 enhances Wnt3a signaling within an optimal concentration range

To investigate the functional roles of the Wnt3a/sFRP2 complex formation, we examined how sFRP2 modulates Wnt3a-mediated signaling by using a TOPFlash reporter system in HEK293T cells (HEK293T/TOPFlash), which monitors the canonical Wnt/β-catenin pathway (**Fig. 4A**).^32^ Incubation of HEK293T/TOPFlash and GFP-Wnt3a with various concentrations of recombinant sFRP2 resulted in a dose-dependent increase of reporter activity up to 400 ng/mL. In contrast, excessive amounts of sFRP2, specifically at 4000 ng/mL in this assay, exhibited no potentiating effect (**Fig. 4B**). We hypothesize that at a condition under excessive amounts of sFRP2, high abundance of free sFRP2 interferes with the transfer of Wnt3a to Fzd receptors. In such environments, the competitive exchange of Wnt3a between free sFRP2 molecules may dominate over receptor engagement.

**Figure 4.**
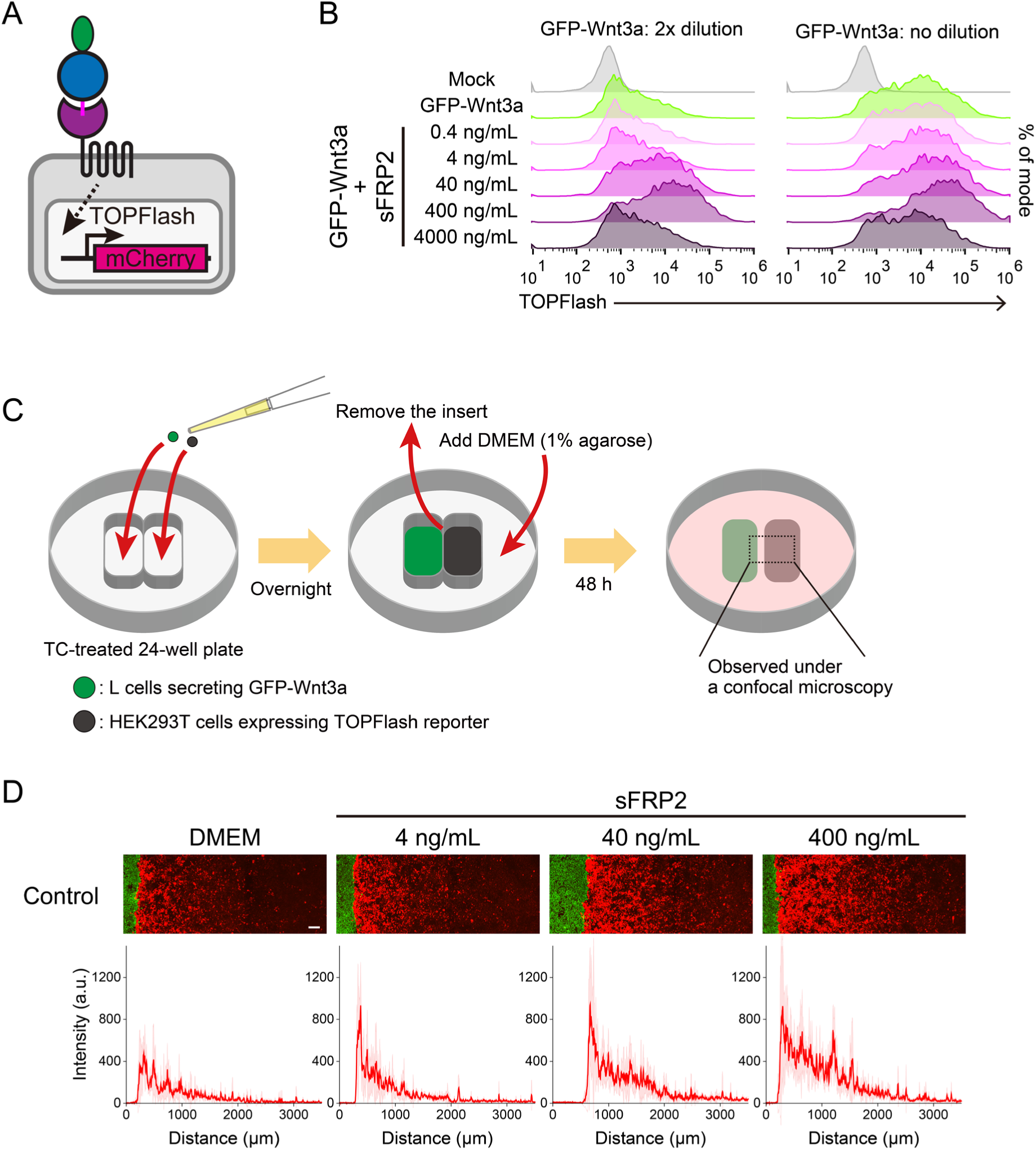
sFRP2 not only enhances canonical signaling pathway but also extends the diffusion length. (A) Schematic illustration of HEK293T cells expressing TOPFlash mCherry reporter. (B) TOPFlash reporter assay with recombinant sFRP2. Reporter cells were incubated with the conditioned media harvested from GFP-Wnt3a-secreting cells and supplemented with various concentrations of recombinant sFRP2. The conditioned media were applied either at full concentration (right) or diluted with an equivalent volume of DMEM (left). After 48-h incubation, the mCherry reporter activation was measured by flow cytometry. The potentiating effect of sFRP2 on Wnt signaling was pronounced under the diluted conditions (left). (C) Experimental setup for the gradient assay. GFP-Wnt3a-secreting L cells (1.6×10^4^ cells) were seeded into the left chamber, while HEK293T TOPFlash reporter cells (1.6×10^4^ cells) were seeded into the right chamber. After incubation overnight, the insert was removed, and the media were replaced with 1% agarose-containing media. Standard DMEM or DMEM containing recombinant sFRP2 was subsequently added over the solidified agarose gel. The confocal images were acquired at 24 h intervals. (D) Gradient patterns after 4 days of incubation. The spatial distribution of mCherry signals was extended in an sFRP2-dose-dependent manner. Red lines indicate the mean mCherry intensity derived from 5 subdivided vertical regions per image. Specifically, each 900-pixel vertical image was partitioned into 5 equal segments of 180 pixels each. The mean fluorescence intensity was calculated for each segment, and the red line represents the average of these five regional means. Shaded areas in the graph represent the ±SD. Scale bar: 200 µm.

Interestingly, the potentiating effect of sFRP2 was more pronounced at lower Wnt3a concentrations (2x dilution of GFP-Wnt3a CM) than at higher concentrations (1x dilution of GFP-Wnt3a CM) (**Fig. 4B**). At saturating Wnt3a concentrations, Wnt3a signaling pathway likely reaches its maximal capacity without sFRP2, masking the contribution of sFRP2. In contrast, under un-saturating Wnt3a concentrations, the pathway remains sensitive to changes in ligand availability. In this regime, sFRP2 effectively transports Wnt3a at the cell surface, thereby increasing receptor engagement and subsequent signal transduction.

We next evaluated the spatial range of Wnt3a signaling activity by establishing a gradient assay using a culture insert system (**Fig. 4C**). GFP-Wnt3a-secreting L cells and HEK293T reporter cells were plated in separate regions, followed by replacement of the medium with 1% agarose-containing medium to prevent convective disturbance.^25^ This gradient assay revealed that higher sFRP2 concentrations significantly extended the signaling range of Wnt3a, indicating that sFRP2 facilitates long-range transport of functional Wnt3a (**Fig. 4D**). In the absence of sFRP2, Wnt3a signaling was restricted to cells most proximal to the source, although Wnt3a is solubilized by forming homo-oligomers or complexes with afamin contained in the culture media. Our observations suggest that such soluble forms of Wnt3a are insufficient for effective membrane recruitment and long-range signaling. Instead, sFRP2 appears to function as a specialized Wnt3a carrier that stabilizes Wnt3a in a soluble yet membrane-associated state, thereby allowing it to navigate the cell surface more efficiently. These mechanisms would be essential for the formation of long-range morphogen gradients during tissue development, where precise spatial control of Wnt signaling is required.

### sFRP2 acts as a Wnt signal enhancer to culture intestinal organoid

To test sFRP2 activity as a Wnt signal enhancer in more physiological contexts such as tissue development, we examined its effect in mouse intestinal organoids. Wnt3a is an essential supplement for various organoid models.^33–35^ Specifically, the Afamin/Wnt3a complex is widely used as a stable and potent supplement, which promotes intestinal organoid growth more effectively than recombinant Wnt3a protein alone.^10^ Prior to the organoid assays, we examined whether sFRP2 enhances Afamin/Wnt3a-mediated signaling using the TOPFlash reporter assay (**Fig. 5A**). Consistent with the results observed with GFP-Wnt3a (**Fig. 4B**), sFRP2 significantly enhanced reporter activity at lower Afamin/Wnt3a concentrations, particularly at 300 ng/mL.

**Figure 5.**
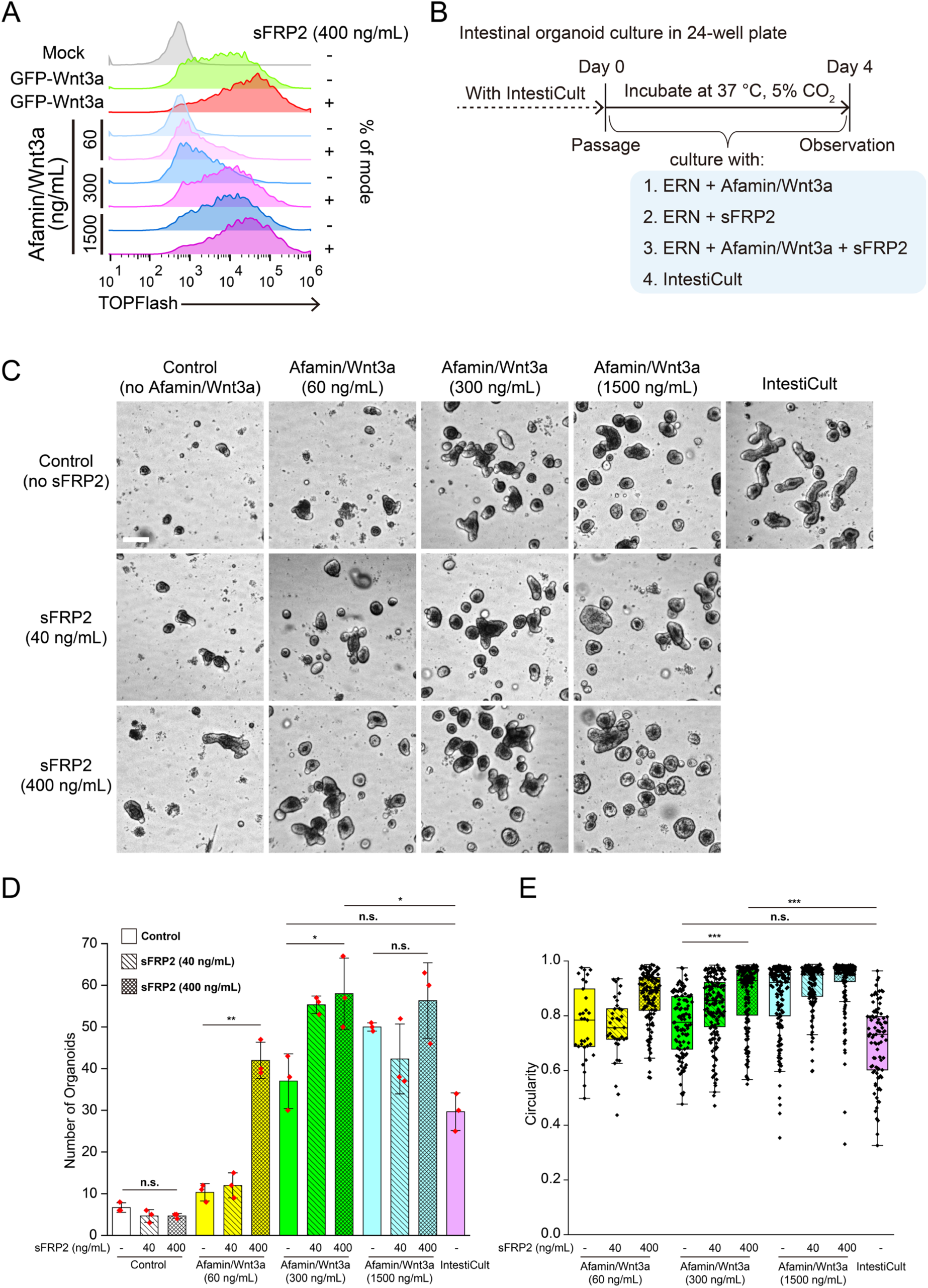
sFRP2 enhances Wnt3a-dependent growth of intestinal organoids. (A) TOPFlash reporter assay with recombinant Afamin/Wnt3a and recombinant sFRP2. The reporter activity was evaluated across a range of recombinant Afamin/Wnt3a concentrations in the presence or absence of 400 ng/mL sFRP2. After 48-h incubation, the mCherry reporter activation was measured by flow cytometry. sFRP2 significantly enhanced reporter activity, with the most significant effect observed at an Afamin/Wnt3a concentration of 300 ng/mL. This synergistic effect is consistent with results obtained from experiments using conditioned media harvested from GFP-Wnt3a-secreting cells. (B) Experimental timeline and organoid culture workflow. When routinely-cultured intestinal organoids with IntestiCult reached passage timing, dissociated organoids were cultured with ERN medium in a 96-well plate. Various concentrations of Afamin/Wnt3a and sFRP2 were supplemented with ERN medium. (C) Representative organoids images with various cultured conditions at 4 days post seeding. Images were acquired every 24 hours by a microscope. Scale bar: 200 µm (D) Quantification of number of expanded organoids. The total number of viable organoids per well was calculated to assess the survival and proliferative impact of each treatment. The addition of sFRP2 led to a significant increase in the number of formed organoids compared to treatments with Wnt3a alone at 60, 300 ng/mL Afamin/Wnt3a concentration. Statistical differences were evaluated using Welch’s t-test with ** for P < 0.01, * for P < 0.05, and n.s. (non-significant) for P ≧ 0.05. Experiments were performed with triplicate, and data are presented as mean ± SD. (E) Quantitative assessment of budding morphogenesis. Organoid circularity was measured to evaluate the shift from spherical cystic structures to complex budding phenotypes. The addition of sFRP2 to Afamin/Wnt3a significantly increased circularity, mirroring the effect of high-dose Wnt3a and suggesting that sFRP2 amplifies Wnt signaling potency. Statistical differences were evaluated using Kruskal-Wallis and Dunn’s post-hoc test with *** for P < 0.001, and n.s. (non-significant) for P ≧ 0.05. Experiments were performed with triplicate, and data are presented as mean ± SD.

Mouse small intestinal organoids were cultured in ERN medium (EGF, R-spondin, and Noggin) supplemented with varying concentrations of Afamin/Wnt3a and sFRP2 (**Fig. 5B**). Brightfield imaging showed that Afamin/Wnt3a promoted both organoid survival and size, whereas sFRP2 alone had a negligible effect (**Fig. 5C**). Strikingly, combined treatment of Afamin/Wnt3a and sFRP2 remarkably enhanced organoid growth, particularly at 60 and 300 ng/mL Afamin/Wnt3a. Quantitative analysis confirmed that 400 ng/mL sFRP2 significantly increased the number of organoids when supplemented with 60 or 300 ng/mL Afamin/Wnt3a (**Fig. 5D**). Furthermore, to evaluate organoid morphology, we analyzed circularity as an indicator of structural maturity. While healthy intestinal organoids typically develop budding crypt structures with low circularity, high Wnt signaling induces a more spherical, cystic morphology with high circularity.^34^ sFRP2 treatment significantly increased organoid circularity, indicating enhanced Wnt activity (**Fig. 5E**). Notably, organoids treated with 60 ng/mL Afamin/Wnt3a plus sFRP2 displayed high circularity comparable to those treated with 1500 ng/mL Afamin/Wnt3a alone. These results demonstrate that sFRP2 effectively amplifies Wnt3a signaling potency. Taken together with the analysis of surface-bound Wnt3a/sFRP2 complexes, our findings suggest that sFRP2 facilitates the recruitment of Wnt3a to the cell surface, thereby achieving robust pathway activation equivalent to that of a much higher ligand dosage.

## Discussion

In this study, we leveraged the mechano-sensing properties of the synNotch receptor to develop a detection system of physical interaction between Wnt3a and the cell surface, which can distinguish soluble and surface-bound forms of Wnt3a complexes. Using this platform, we demonstrated that sFRP2 facilitates the recruitment of Wnt3a complexes to the cell surface and their subsequent internalization. Beyond the specific interactions examined here, our work highlights the broader utility of synNotch receptors as a powerful platform for investigating the extracellular dynamics of various surface-associated signaling molecules. This approach can be readily extended to study other lipid-modified morphogens, such as Sonic hedgehog (Shh) and Bone Morphogenetic Proteins (BMPs),^36,37^ whose solubilization, transport, and presentation are similarly regulated by complex interactions with HSPGs and other membrane-associated factors.

Our findings reveal that sFRP2 functions as an extracellular modulator of Wnt3a signaling rather than as an antagonist of Wnt3a, as traditionally proposed,^38^ suggesting that sFRP2 may have multiple roles in different tissue contexts. In our *in vitro* mammalian cell culture system, the association between Wnt3a and sFRP2 not only enhances canonical Wnt/β-catenin pathway but also extends the effective signaling range of Wnt3a. The results obtained using intestinal organoid cultures further underscore the physiological relevance of this mechanism. In this system, sFRP2 acts as a potent enhancer that compensates for limited Wnt3a availability, shifting organoid morphology toward a spherical structure that is otherwise observed only at much higher ligand concentrations. This capacity to modulate both signaling strength and spatial distribution of Wnt3a implies sFRP2 as a critical regulator in the formation and maintenance of Wnt morphogen gradients.

Direct visualization of Wnt3a transport to its receptor will be essential to further substantiate the molecular mechanisms proposed here. High-speed atomic force microscopy (HS-AFM) offers a promising approach, as it enables real-time observation of individual molecules in aqueous environments.^39,40^ In particular, HS-AFM could be utilized to visualize dynamic molecular exchange between Wnt3a/sFRP2 complexes and Wnt3a/Frizzled receptor complexes. Moreover, this approach using recombinant Wnt3a/sFRP2 complexes would provide definitive evidence for direct binding interactions between sFRP2 and HSPGs at the single-molecule level.

An important remaining question is whether Wnt3a/sFRP2 complexes elicit physiological effects beyond activation of the canonical Wnt pathway. From the perspective of physiological roles in vivo, analyses of sFRP2 knockout mice have revealed defects in bone regeneration and reduced Wnt signaling in skeletal stem cells, supporting the notion that sFRP2 can function as a Wnt agonist in physiological contexts.^41^ It will therefore be interesting to examine how sFRP2 regulates Wnt signaling during tissue development and regeneration in vivo using sFRP2-deficient models. In parallel, precise control of Wnt activity is critical in vitro for regulating proliferation and differentiation of stem cells within diverse organoid systems.^42^ Comprehensive transcriptomic profiling and gene expression analyses of organoids cultured with or without sFRP2 will be essential to elucidate additional roles of sFRP2 in shaping Wnt-dependent cell fate decisions. More broadly, strategies that regulate Wnt activity and gradient formation through extracellular carrier proteins may provide a rational framework for optimizing and engineering organoid culture.

## Methods

### Plasmid construct and cell culture

Plasmid constructs: mouse Fzd8, mouse sFRP2, mouse sFRP3, and L cells expressing FLAG-tagged mouse Wnt3a were kindly gifted by Prof. Shinji Takada (National Institutes of Natural Sciences). HepIII-HA-GPI (membrane-tethered HepIII) was kindly gifted by Dr. Takafumi Ikeda and Masanori Taira. All provided plasmid constructs are translocated to lentiviral plasmid vector, pHRSIN:CSW, with puroR gene transcribed by PGK promoter to undergo drug selection and establish stable cell lines. To establish chimeric sFRP2s, GFP or mCherry were directly fused to the C-terminus of corresponding sFRP2 domains (**Fig. 3D and Fig. S1A**). TM-CRD consists of a CRD domain of sFRP2 and a PDGFR transmembrane domain.

Following the procedure we previously reported, we generated detector cell lines expressing anti-GFP LaG17 or LaG2 synNotch receptor (Addgene, #162230), together with Tetracycline Response Element (TRE)-based reporters (Addgene, #162231). TOPFlash-mCherry reporter was optimized from TOPFlash-GFP reporter (Addgene, #35489).

All cell lines, L cells, L929 cells (RIKEN BRC, RCB1422), MDCKII cells (RIKEN BRC, RCB5148), EpH4 cells gifted by Prof. Yasuyuki Fujita, NIH3T3 cells (gifted by Prof. Rikinari Hanayama), HEK293T cells (RIKEN BRC, RCB2202) were maintained at 37 °C in a humidified 5% CO_2_ and cultured in DMEM (Nacalai Tesque, #08458-16) containing 10% fetal bovine serum (FBS) (Gibco) and 1x penicillin-streptomycin (Wako, #168-23191). Cells were usually grown on 60-mm tissue culture-treated dishes (Greiner, #628160), whereas protein-secreted cells such as GFP-Wnt3a and sFRP2 were cultured in 100-mm tissue culture-treated dishes (Greiner, #664160) to obtain conditioned media.

### Lentiviral transduction

All stable cells in this study were established using lentiviral transduction. For virus production, HEK293T cells were transfected with the target plasmid vector, pCMVdR8.91 and pMD2.G using PEI MAX (Polysciences, #24765). To generate the stable lines, 0.8×10^5^ L929 cells were seeded into 12-well plates (Corning, #3513) and incubated with varying volumes of the lentiviral supernatant in a DMEM-based solution containing 10 μg/mL hexadimethrine bromide (Sigma-Aldrich, #H9268).

### Preparation of conditioned media

Cells were maintained in 100 mm tissue culture-treated dishes containing 10 mL of DMEM supplemented with 10% FBS. Upon reaching 100% confluency, the cultures were incubated for an additional 24 hours. The resulting supernatants were collected using a 10 mL syringe (Terumo, #SS-10SZ) and subsequently passed through a 0.45 µm filter (Merck, #SLHVR33RB). The biological activity of proteins in conditioned media remained stable for at least two weeks when stored at 4 °C.

### Assays for ligand detection using synNotch cells and TOPflash assay

To detect surface-bound ligands, 2.0×10^4^ detector cells were seeded into each well of a 96-well plate (Corning, #3595) and maintained in 200 µL of DMEM-based conditioned media. After 24-hour incubation, the activation levels of the detector cells were quantified by flow cytometry (Beckman Coulter, CytoFLEX S). Fluorescence images were acquired using confocal microscopy (ANDOR, DragonFly or Nikon, AX). For co-culture experiments involving membrane-tethered ligands, such as mFzd1, mFzd8, and TMD-CRD (**Fig. 1D and 2D**), 1.0×10^4^ detector cells were mixed with an equal number of cells expressing membrane-tethered ligands. These cell mixtures were then processed according to the same protocol described above. For the activation assay in a 3D culture environment (**Fig. S1B-C**), 150 detector cells were seeded into a 384-well round-bottom ultra-low-attachment plate (Corning, #3830) with conditioned media. After 24-h incubation, the fluorescent levels were observed under confocal microscopy (Nikon, AX).

To evaluate the activity of Wnt3a signaling, 1.0×10^4^ HEK293T expressing TOPFlash-mCherry reporter were plated into each well of a 96-well plate with 200 µL of DMEM-based conditioned media. After 48-hour incubation, the signaling activation was analyzed by flow cytometry.

The acquired flow cytometry data were analyzed using the FlowJo software. Confocal images were analyzed using ImageJ software.

### Gradient assay using culture insert

To establish a stable GFP-Wnt3a or Wnt3a gradient, a two-well culture insert (ibidi, #80209) was centered within each well of a 24-well plate. The left chamber of the insert was populated with 1.6×10^4^ Wnt3a-secreting cells in 80 µL of medium, while the same number of TOPFlash reporter cells were added to the right chamber with an equivalent volume. The surrounding area outside the insert was filled with 300 µL of DMEM. After a gentle centrifugation at 100 x g for 2 minutes, the plate was incubated overnight. After the incubation, the left chamber was gently rinsed twice with 80 µL of medium. The insert was removed and supernatant was aspirated, then 800 µL of 1% Agarose L (NIPPON GENE, #317-01182) was added. This agarose medium was prepared by mixing 2% agarose in PBS (Wako, #049-29793) with 20% FBS-containing DMEM at a 1:1 ratio. After the gel was solidified at room temperature for 15 minutes, 1 mL of sFRP2 conditioned medium or DMEM as control was added on the agarose medium. The images were obtained every 24 hours for 5 days by confocal microscopy (Nikon, AX).

### Intestinal organoid

Mouse small intestinal organoids were maintained inside 50% Matrigel (Corning) in IntestiCult Organoid Growth Medium according to the manufacture’s protocol (VERITAS, #ST-06005). For experimental assays, organoids were dissociated into single cells or small clusters by Cell Recovery Solution (Corning, #354253). Dissociated organoids with 50% Matrigel were seeded into a 96-well plate (Corning, #353219) in defined ENR medium. This medium consisted of Advanced DMEM/F12 supplemented with 10 mM HEPES (Gibco, #15630106), 1 x GlutaMAX (Gibco, #35050061), 1 x N2 (Gibco, #17502048), 1 x B27 (Gibco, #17504044), 1 mM N-acetylcysteine (Sigma, A9165-5G), 100 ng/mL recombinant murine EGF (Gibco, #PMG8041), 100 ng/mL recombinant murine Noggin (Peprotech, #250-38-20UG), 50 ng/mL recombinant murine R-spondin-1 (R&D Systems, 3474-RS-050), and 10 µM Y27632 (Wako, #030-24021).

Organoid phenotypes were evaluated by supplementing the ENR medium with recombinant Afamin/Wnt3a and mouse sFRP2 (R&D Systems, #1169-FR-025) at the indicated concentrations. The data were acquired four days after tissue culture by monitoring organoid morphology using fluorescence microscopy (Keyence, BZ-X800).

### Image analysis

To quantify the internalization of Wnt3a (**Fig. 3C-D and Fig. S4A**), each single cell was manually segmented based on differential interference contrast (DIC) images. During this segmentation process, overlapping cells were specifically excluded to ensure accurate measurements. The total fluorescence intensity for each individual cell was subsequently analyzed using ImageJ software. The degree of colocalization between GFP-Wnt3a and mCherry-sFRP2 was assessed by calculating Pearson’s correlation coefficients using the JaCoP plugin in ImageJ, after performing the automatic segmentation using Cellpose2.0 (**Fig. 3D and Fig. S5**).^43^ To eliminate the possibility of coincidental colocalization, the correlation values obtained from the original dual-channel images (GFP and mCherry) were compared against those derived from randomly shuffled GFP and mCherry channels. This statistical validation was performed according to Costes’ automatic thresholding and randomization method. In **Fig. 5C-E**, individual organoids were automatically segmented from brightfield images using Cellpose 2.0. From the resulting masks, the total organoid count and circularity were quantified using ImageJ software.

## Acknowledgments

We thank S. Takada, R. Takada, Y. Mii, T. Ikeda and M. Taira for providing materials and for insightful discussions. We also thank members of the Toda laboratory for discussion and assistance.

## Funding

This work was supported by the Japan Science and Technology Agency (JST), PRESTO Grant No. JPMJPR2147 and FOREST Grant No. JPMJFR2311; the Japan Society for the Promotion of Science (JSPS), KAKENHI Grant No.21H05291 and 24K02021, Japan; Joint Research of the Exploratory Research Center on Life and Living Systems (ExCELLS) (ExCELLS program No. 22EXC205 and 23EXC203), Japan to S.T.; and World Premier International Research Center Initiative (WPI), Ministry of Education, Culture, Sports, Science and Technology (MEXT), Japan to K.M. and S.T.; K.M. is supported by the Yoshida Scholarship Foundation PhD fellowship and the ANRI fellowship.

## Author contributions

K.M. developed and planned research; carried out design, construction, and performed experiments; and wrote and edited the manuscript. S.T. developed, planned, and oversaw research; wrote and edited the manuscript.

## Competing interests

S.T. is an inventor on a patent for synthetic Notch receptors (Patent No.: US 10,590,182 B2) held by the Regents of the University of California, which is licensed to Gilead. K.M. declares no competing interests.

## Data and materials availability

All data are available in the main manuscript and supplementary materials.

