## Supplementary information for "A synNotch-based morphogen detection system reveals sFRP2 enhances Wnt3a signaling"

Kosuke Mizuno *et al.*

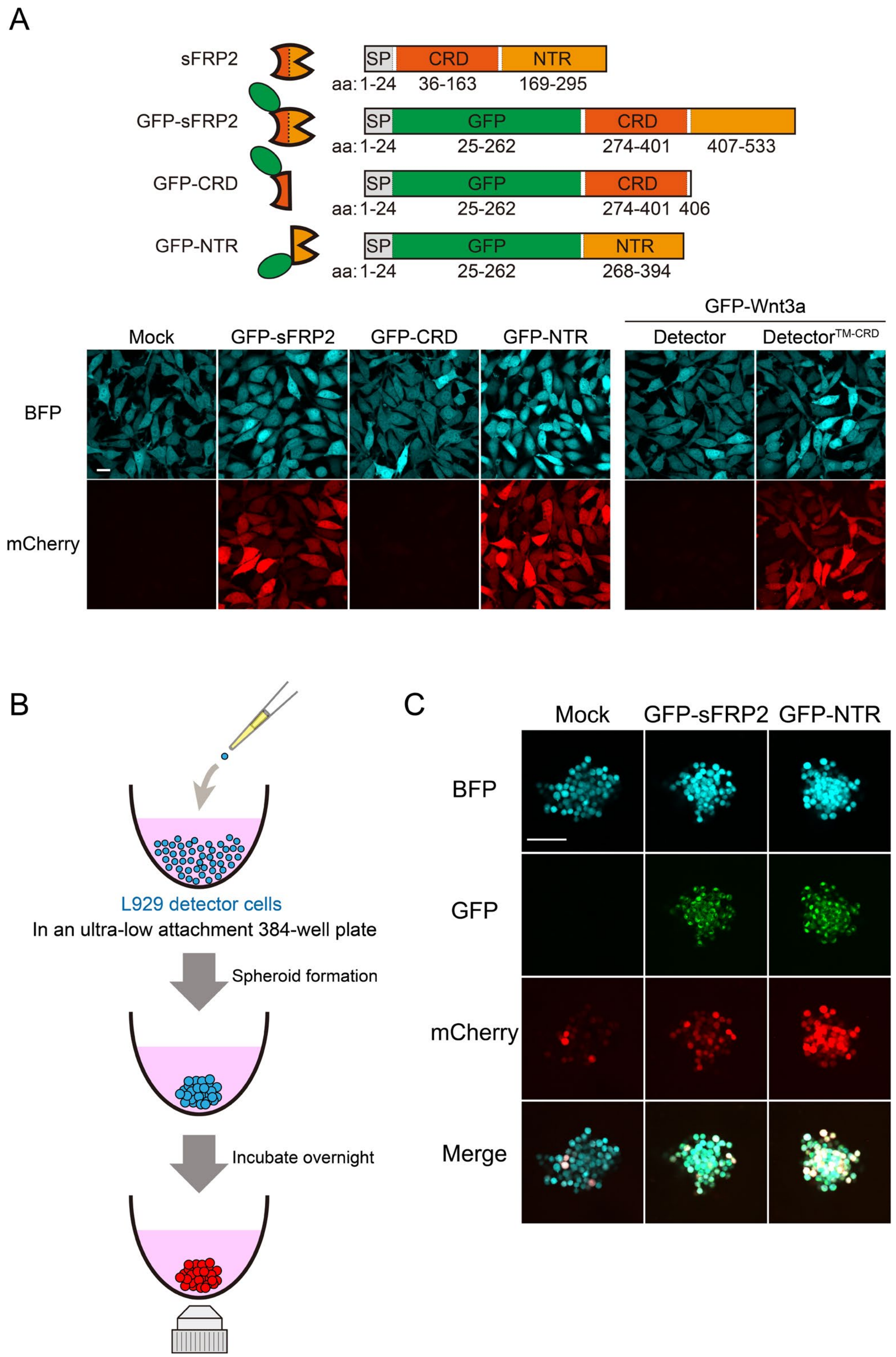

**Supplementary Fig. S1. Activation of detector cells in a 3D culture environment**

**(A)** Construct schematics of chimeric sFRP2s and confocal images of detector cells treated with chimeric sFRP2s in **Fig. 1D**. The mCherry reporter was induced by treatment with either GFP-sFRP2, GFP-NTR, or GFP-Wnt3a in the presence of TM-CRD. tagBFP is stably expressed in detector cells as a marker for reporter cassette integration. Scale bar: 20 µm. **(B)** Schematic illustration of the 3D assay. Spheroids consisting of 150 detector cells were formed in an ultra-low attachment 384-well plate with conditioned media. After overnight incubation, fluorescent signals were observed using confocal microscopy. **(C)** Confocal images of detector spheroids. The mCherry signals observed in mock treatment indicated basal leakage. Coculture with GFP-sFRP2 or GFP-NTR led to significant activation even in a 3D environment. Scale bar: 100 µm.

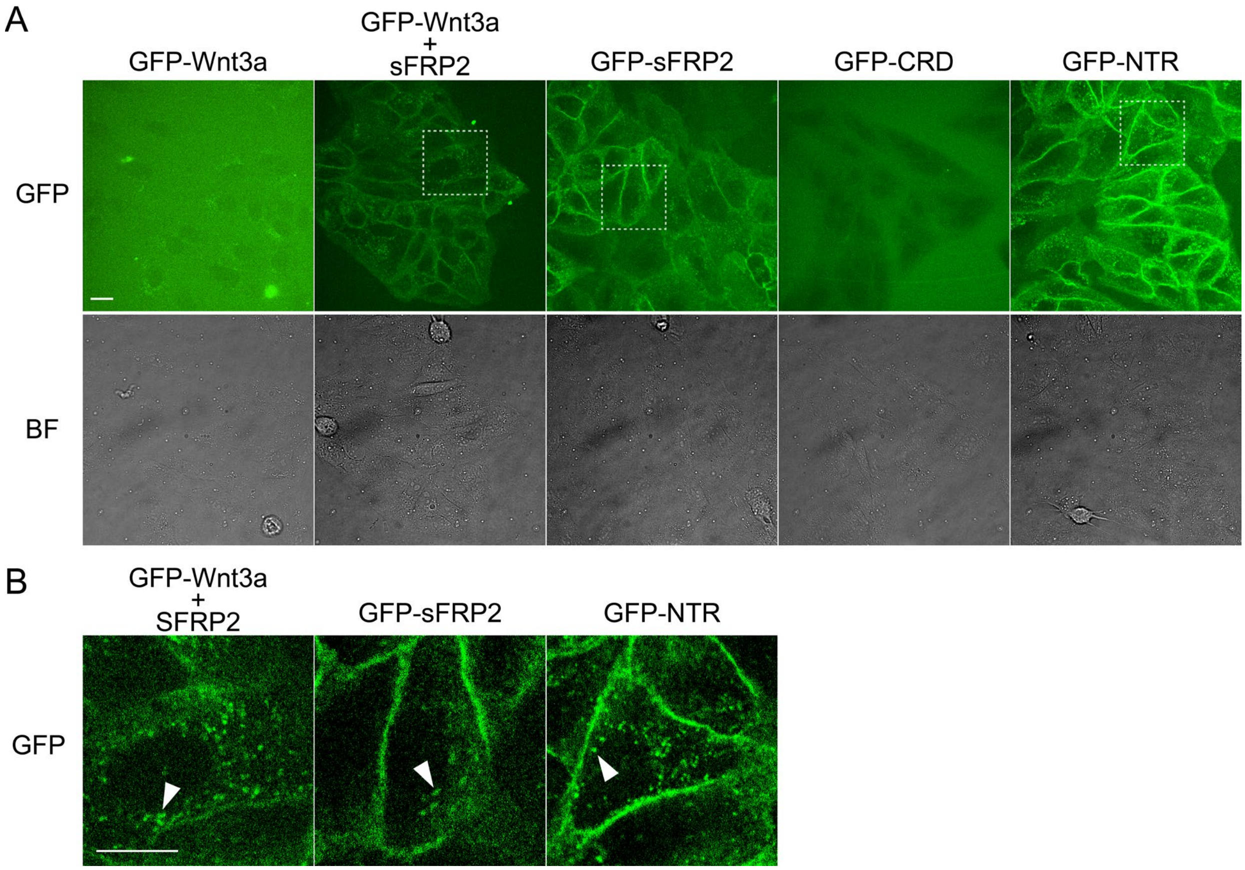

**Supplementary Fig. S2. Localization of GFP-Wnt3a and chimeric sFRP2**

**(A)** Confocal images of wild-type MDCKII cells treated with conditioned media. The combined treatment of GFP-Wnt3a and sFRP2, as well as GFP-sFRP2 and GFP-NTR alone, showed localization of GFP signals near the cell surface, whereas GFP-Wnt3a alone showed no accumulation. Scale bar: 20 µm. **(B)** Magnified images of dashed squares shown in (A). White arrow heads indicate internalized puncta. Scale bar: 20 µm.

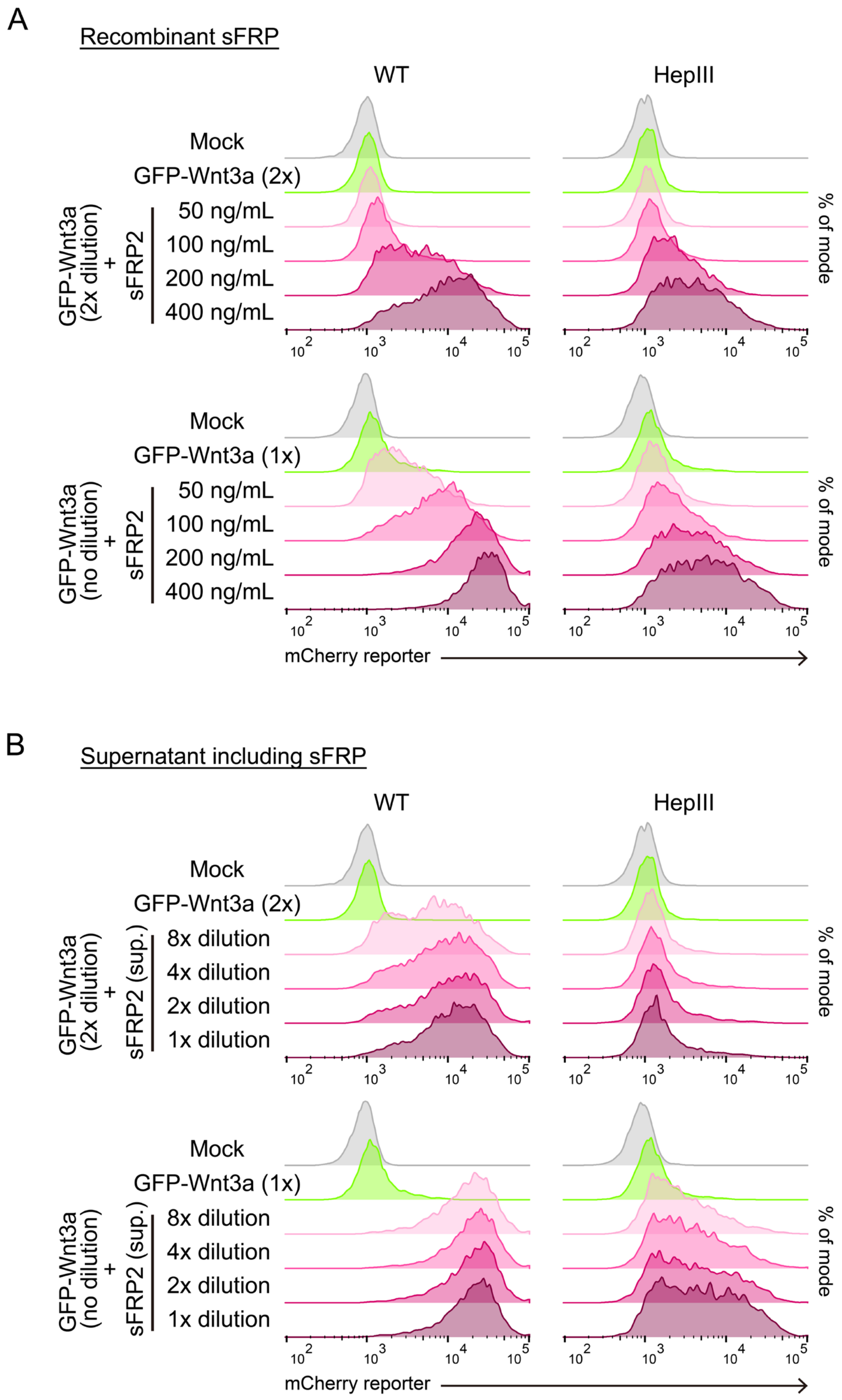

**Supplementary Fig. S3. sFRP2 dose-dependent activation of L929 detector cells**

**(A)** Flow cytometry analysis of L929 detector cells treated with GFP-Wnt3a and recombinant sFRP2. Detector cells were activated by GFP-Wnt3a in a sFRP2 concentration-dependent manner, which was inhibited with HepIII. 2x dilution of GFP-Wnt3a concentration reduced activation levels while maintaining the trend. (**B**) Flow cytometry analysis using sFRP2 CM. When the media was diluted up to 8-fold, the activation level with 2x dilution GFP-Wnt3a CM showed a level at between 200 ng/mL and 400 ng/mL of recombinant sFRP2 treatment in (A). This suggests that the conditioned media contained concentrations between 200 ng/mL and 400 ng/mL of sFRP2.

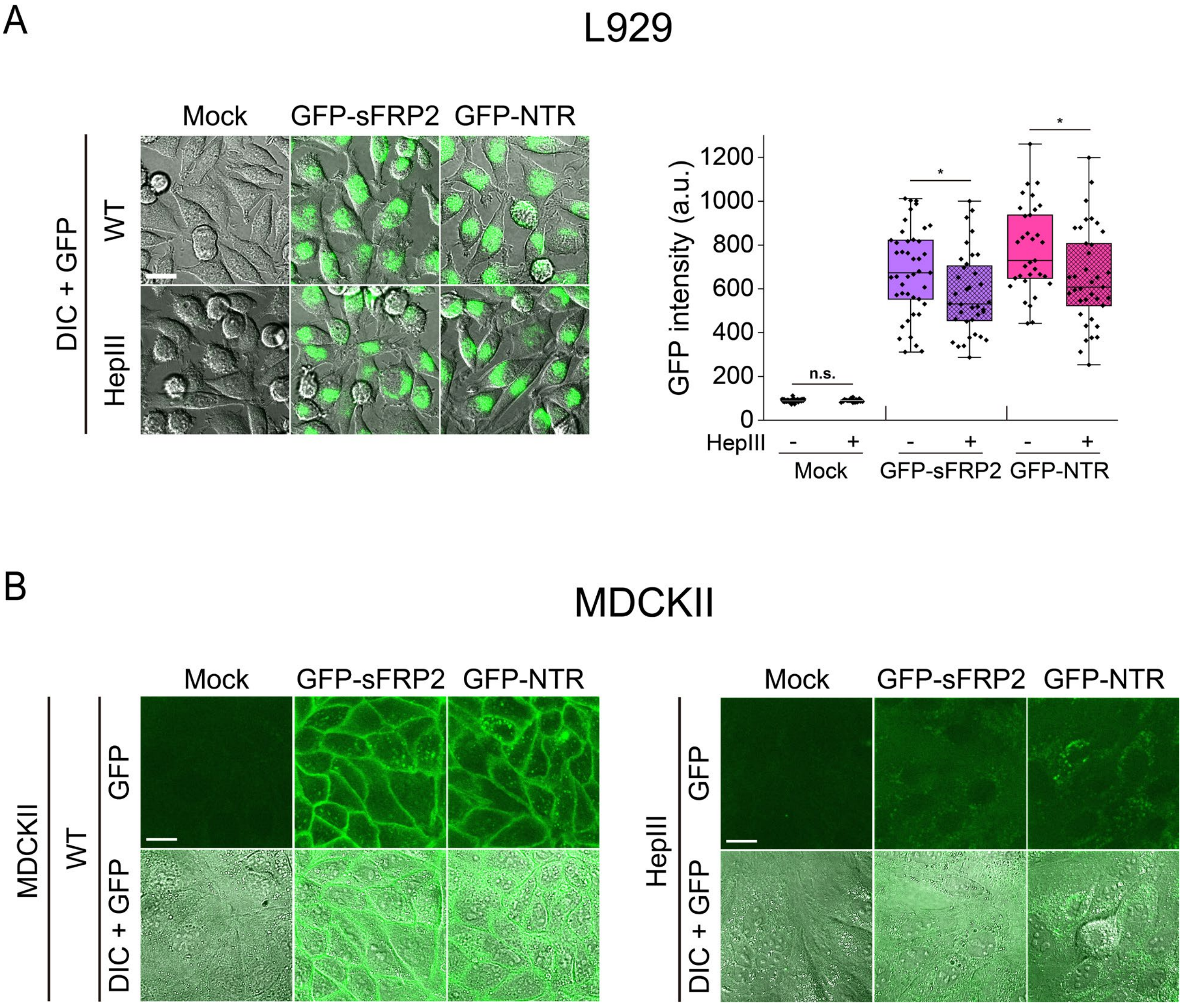

**Supplementary Fig. S4. Heparan sulfate-dependent internalization of Wnt3a and sFRP2**

**(A)** Quantification of chimeric sFRP2 recruitment and internalization. After 24 hours of treatment with conditioned media, confocal images were acquired for the GFP channel and differential interference contrast (DIC) (left). The total GFP intensity in each single cell was quantified after performing segmentation (right). Scale bar: 20 μm. Treatment with GFP-sFRP2 and GFP-NTR both led to increased GFP signals relative to the mock treatment. HepIII-expressing cells exhibited a modest reduction in GFP recruitment under both conditions, yet substantial GFP signals persisted within the intracellular space. Statistical differences were evaluated using Welch's t-test with * for $P < 0.05$, and n.s. (non-significant) for $P \geqq 0.05$. **(B)** Confocal images of wild-type and HepIII-expressing MDCKII cells treated with conditioned media. GFP accumulation on cell surface was significantly inhibited by HepIII when cultured with conditioned media of GFP-sFRP2 or GFP-NTR. Scale bar: 20 μm.

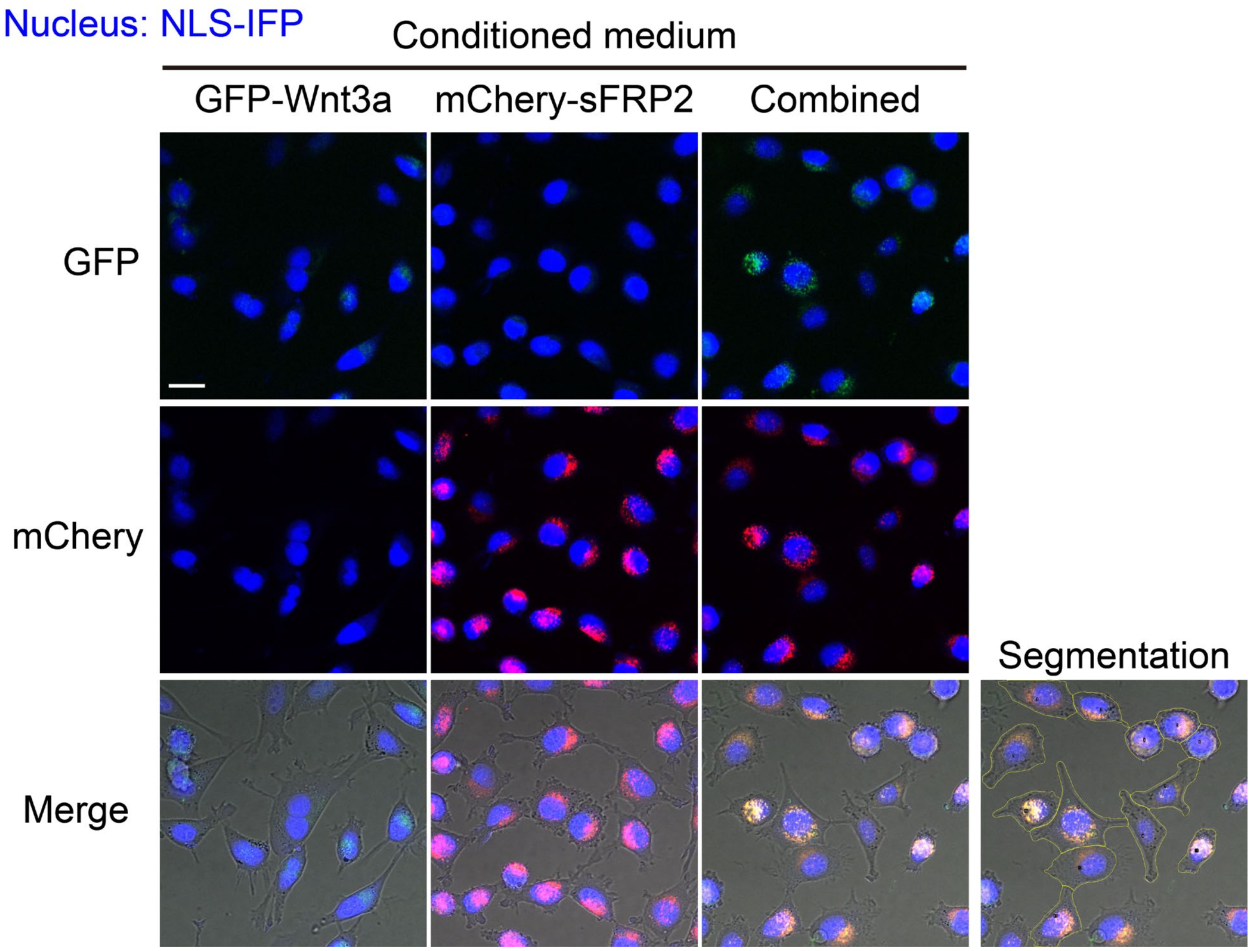

**Supplementary Fig. S5. Co-localization of GFP-Wnt3a and mCherry-sFRP2**

Confocal images of wild-type L929 cells treated with GFP-Wnt3a and mCherry-sFRP2. Treatment with conditioned media containing GFP-Wnt3a alone showed no accumulation of GFP inside the cells, whereas combined treatment with mCherry-sFRP2 induced GFP internalization. The rightmost image shows the result after segmentation processing, indicated by yellow lines. Scale bar: 20 µm.
